# Polygenic Prediction of Substance Use Disorders in Clinical and Population Samples

**DOI:** 10.1101/748038

**Authors:** Peter B. Barr, Albert Ksinan, Jinni Su, Emma C. Johnson, Jacquelyn L. Meyers, Leah Wetherill, Antti Latvala, Fazil Aliev, Grace Chan, Samuel Kuperman, John Nurnberger, Chella Kamarajan, Andrey Anokhin, Arpana Agrawal, Richard J. Rose, Howard J. Edenberg, Marc Schuckit, Jaakko Kaprio, Danielle M. Dick

## Abstract

Genome-wide, polygenic risk scores (PRS) have emerged as a useful way to characterize genetic liability using genotypic data. There is growing evidence that PRS may prove useful to identify those at increased risk for developing certain diseases. The current utility of PRS in relation to alcohol use disorders (AUD) remains an open question. Using data from both a population-based sample [the FinnTwin12 (FT12) study] and a high risk sample [the Collaborative Study on the Genetics of Alcoholism (COGA)], we examined the association between PRSs derived from genome-wide association studies (GWASs) of 1) alcohol dependence/alcohol problems, 2) alcohol consumption, and 3) risky behaviors with AUD and other substance use disorder (SUD) symptoms. Individuals in the top 20%, 10%, and 5% of PRSs had increasingly greater odds of having an AUD compared to the lower end of the continuum in both COGA (80^th^ % OR = 1.95; 90^th^ % OR = 2.03; 95^th^ % OR = 2.13) and FT12 (80^th^ % OR = 1.77; 90^th^ % OR = 2.27; 95^th^ % OR = 2.39). Those in the top 5% reported greater levels of licit (alcohol and nicotine) and illicit (cannabis) SUD symptoms. PRSs can predict elevated risk for SUD in independent samples. However, clinical utility of these scores in their current form is modest. As these scores become more predictive of SUD, they may become useful to practitioners. Improvement in predictive ability will likely be dependent on increasing the size of well-phenotyped discovery samples.

## Introduction

Alcohol misuse is one of the leading contributors to preventable mortality and morbidity worldwide ^1-3^. Identifying individuals at heightened risk for developing alcohol related problems remains an important goal of medical practitioners. One important risk factor for alcohol misuse is one’s own genetic liability. Twin and family studies indicate that genetic influences on alcohol use disorders (AUD) account for approximately 50% of the variation in the population ^4^. Genome-wide association studies (GWASs) have identified multiple variants associated with AUD ^5-7^, alcohol consumption ^7, 8^, and maximum alcohol intake^9^. Using information from these GWASs, we are now able to aggregate risk across the genome by creating genome-wide polygenic scores (PRS) to predict risk in independent samples ^5, 6, 8, 10^.

Beyond being useful for research purposes, researchers have begun to examine the clinical utility of PRS to predict risk for medical outcomes. PRS for coronary artery disease (CAD), atrial fibrillation (AF), type 2 diabetes (T2D), inflammatory bowel disease (IBS), and breast cancer (BC) have been found to be as predictive of these diseases as well known monogenic mutations ^11^. Individuals in the top 5% of the PRS distributions had ∼3 fold likelihood of having CAD, AF, T2D, IBS, or BC compared to the bottom 95%. For obesity, individuals in the top PRS decile were on average 13 kg heavier than those in the bottom decile ^12^. These studies demonstrate the potential for identifying individuals at heightened risk for various medical conditions using PRS. However, the clinical utility of PRSs for AUD in relation to substance use phenotypes remains an open question.

In the current analysis, we tested PRS in two target samples, a population-based sample and a clinically ascertained sample of families deeply affected by AUD, to evaluate the current state of alcohol-related PRS for possible clinical utility. We use several discovery samples from large scale GWAS to create three PRS: a meta-analysis of two GWASs on alcohol-related problems ^5, 6^, a recent large-scale GWAS of alcohol consumption ^8^, and a GWAS for risky behaviors, including alcohol use ^13^. We chose to test PRS based on multiple alcohol-related GWAS because multiple lines of evidence indicate alcohol consumption and dependence have only partially shared genetic etiology ^5, 6, 14, 15^. Additionally, we include a PRS for general risk behavior as there is robust evidence that the genetic risk for alcohol and other substance use disorders is shared with other disorders and behaviors related to reduced inhibitory control. This constellation of behaviors is often referred to as the externalizing spectrum ^16-18^. We test the association of these PRS with a variety of substance use outcomes (including alcohol, nicotine, and other illicit substance use disorders), based on the robust finding that substance use disorders share an underlying genetic architecture, with the majority of the heritability shared across substances ^16-18^.

## Methods

### Samples

*The FinnTwin12 Study (FT12)* is a population-based study of Finnish twins born 1983-1987 identified through Finland’s Central Population Registry. A total of 2,705 families (87% of all identified) returned the initial family questionnaire late in the year in which twins reached age 11. Twins were invited to participate in follow-up surveys when they were ages 14, 17, and approximately 22 (during young adulthood). An intensively studies sample was selected as 1035 families, among whom 1854 twins were interviewed at age 14. The interviewed twins were invited as young adults to complete the Semi-Structured Assessment for the Genetics of Alcoholism (SSAGA) ^19^ interview (n = 1,347) and provide DNA samples (see Kaprio 2013 for a full description). The current analysis uses data from the young adult wave (mean age = 21.9; range 20-26), which included retrospective lifetime diagnoses.

*The Collaborative Study on the Genetics of Alcoholism (COGA)* is a sample of high-risk families ascertained through adult probands in treatment for AUD and a smaller set of comparison families from the same communities. In the first 10 years, probands along with all willing first-degree relatives were assessed; recruitment was extended to include additional relatives. Data collection included the SSAGA 19, neurophysiological and neuropsychological protocols, and collection of blood for DNA. In 2004, COGA began a prospective study of adolescents and young adults, targeting assessment of youth aged 12-22 from COGA families where at least one parent had been interviewed. These young participants were re-assessed every two years. The sample is racially/ethnically diverse (60.6% non-Hispanic White, 24.9% Black, 11.1% Hispanic, and 3.4% other). Most (84%) have GWAS data. A full description of the COGA sample is available elsewhere ^21-23^. For the present study, we only focused on COGA participants of empirically assigned (as verified from GWAS data) European ancestry (n = 7,599) because each of the discovery GWAS samples were primarily of European ancestry. Ancestral mismatch between discovery and target samples can lead to bias in the performance of polygenic scores ^24^.

### Measures

#### Alcohol Use Disorder (AUD)

We used SSAGA interviews to construct lifetime symptom counts of DSM-5 AUD ^25^ in each sample. Because individuals in COGA are potentially interviewed multiple times, we used the highest symptom count ever reported by each subject. In FT12, lifetime symptom counts were measured at the young adult interview. In addition to symptom counts, we created AUD thresholds for those who met criteria for mild (2+ symptoms), moderate (4+ symptoms), or severe (6+ symptoms) AUD ^25^ without clustering. In both FT12 and COGA, individuals who had never initiated alcohol use were coded as missing.

#### Other Substance Use Disorders (SUD)

We constructed lifetime symptom counts of cannabis, cocaine, and opioid use disorders based on DSM-5 criteria. We measured nicotine dependence symptoms using the Fagerstrom Test for Nicotine Dependence (FTND), which assesses six symptoms and has values ranging from 0 to 10 in both COGA and FT12. Because many illicit SUDs were not measured or rare in the FT12 data, we limit analyses of illicit SUD to COGA. Like AUD, these symptom counts represent the maximum reported for each respondent across the course of the study. Symptom counts for each substance were limited to those who indicated ever using the corresponding substance. In the case of FTND, this is limited to those who report smoking 100+ cigarettes in their lifetime.

#### Polygenic Scores (PRS)

We created PRS derived from publicly available large-scale GWASs. Information on genotyping and quality control is available in the supplemental information. We used the well-established process of clumping and thresholding ^26^. Single nucleotide polymorphisms (SNPs) from discovery GWASs were clumped based on linkage disequilibrium (LD) in the 1000 genomes EUR panel using PLINK ^27^, based on an R^2^ = .25, with a 500 kb window. SNPs were weighted using the negative log of the association p-values. We then created scores based on differing thresholds of GWAS p-values (p<.0001, p<.001, p<.01, p<.05, p<.10, p<.20, p<.30, p<.40, p<.50). We converted PRS to Z-scores for interpretation.

We used four primary discovery GWASs to create three different PRSs. The first was from a recent GWAS of number of alcoholic drinks per week in approximately one million individuals provided by the GWAS & Sequencing Consortium of Alcohol and Nicotine Use (GSCAN) ^8^. We obtained GSCAN summary statistics with all Finnish (which included FinnTwin12) and 23andMe (which are not publicly available) cohorts removed (available N = 534,683). The PRS for alcohol problems were derived from a meta-analysis of two GWASs: a GWAS on the problem subscale from the Alcohol Use Disorders Identification Test (questions 4-10; AUDIT-P) in 121,604 individuals from the UK Biobank ^6^ and the Psychiatric Genomcs Consurtium’s (PGC) GWAS of alcohol dependence (N = 46,568) ^5^. Both FT12 and COGA were in the initial AD GWAS and we obtained summary statistics with each cohort removed (meta-analysis results available in supplemental info). Finally, we derived a PRS for risky behaviors from a GWAS of the first prinicipal component of four risky behaviors (drinks per week, ever smoking, propensity for driving over the speed limit, and number of sexual partners) from 315,894 individuals in the UK Biobank ^13^. While this PRS does include alcohol consumption and smoking, it captures the shared variance between these substance use measures and the other two risky behaviors. These polygenic scores covered the domains of alcohol consumption (GSCAN DPW), alcohol problems (PROB ALC), and general externalizing (RISK PC).

#### Analytic Strategy

We identified the most predictive PRS across p-value threshold from each of the discovery GWASs in both COGA and FT12 using the change in R^2^ above a baseline model with sex, age of last observation, the first ten ancestral principal components (PCs), genotyping array, and data collection site (these latter two were only included in COGA analyses). We used linear/generalized-linear mixed-effects models with random intercepts to adjust for clustering at the family level and a pseudo-R^2^ for mixed models ^28^. After identifying the most predictive PRS, we estimated the joint effect of all PRS on AUD symptoms to examine whether each PRS explained unique variance. We next divided PRSs at various thresholds (80^th^, 90^th^, and 95^th^ percentiles) to determine the increase in likelihood of AUD (using symptom severity thresholds of AUD) associated with being in the top end of split relative to the bottom portion of the split. Because increased risk ratios do not necessarily reflect clinical utility ^29^, we also calculated area under the curve (AUC) of the joint model containing all continuous PRS to estimate sensitivity/specificity (in supplemental information). Finally, we compared mean values of other substance use outcomes for the top 5% in each PRS to those in the bottom 95%.

## Results

Table 1 presents the descriptive statistics for each of the samples. Each sample has slightly more female than male participants. COGA has a broader age range and higher mean age. As COGA was primarily ascertained for families with multiple AUD members, the mean number of AUD symptoms (mean = 3.44) is significantly higher than in the population-based FT12 sample (mean = 1.63). Additionally, COGA participants had higher mean levels of FTND symptoms (mean = 4.17) than FT12 participants (mean = 2.57). For other SUD symptoms in COGA, though symptom counts for cannabis, cocaine, and opioid use disorders are zero-inflated; there are a substantial number of participants who report non-zero levels of symptoms (see Table 1).

**Table 1:**
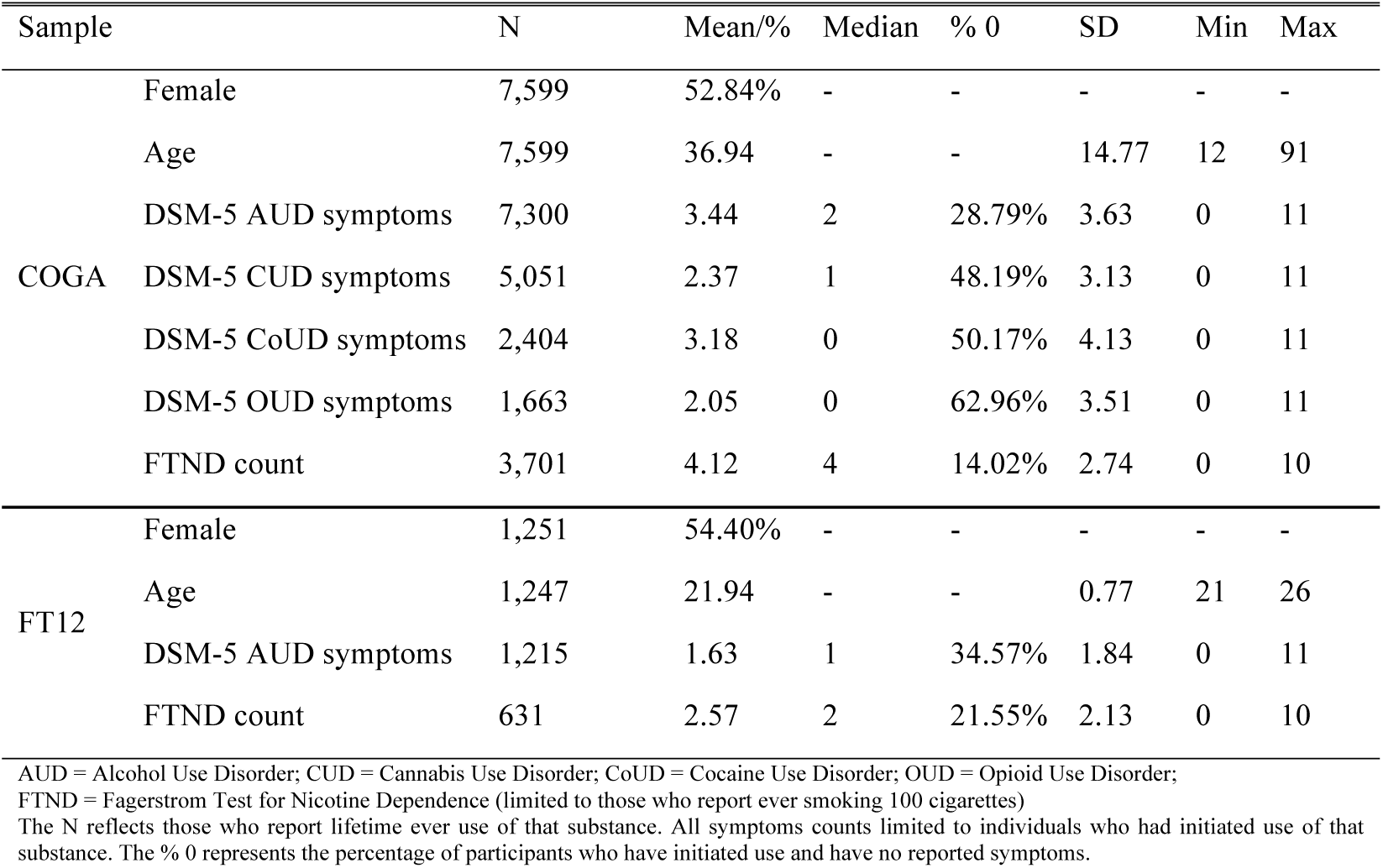
Descriptive Statistics for FT12 and COGA samples.

### Predictive Power of PRS

Figure 1 presents the ΔR^2^ using each PRS to predict AUD symptom count, in each sample, for each discovery GWAS p-value threshold. Within each sample, we chose the most predictive PRS for each discovery GWAS in COGA (RISK PC = p <.10, GSCAN DPW = p <.05, PROB ALC = p <.50) and FT12 (RISK PC = p <.10, GSCAN DPW = p <.20, PROB ALC = p <.50). In COGA, the PRS for GSCAN DPW was the most predictive of AUD symptoms (ΔR^2^ = 1.80%). The RISK PC PRS was most predictive of AUD symptom count in FT12 (ΔR^2^ = 2.10%). These PRS were followed by PRS for risky behaviors in COGA (ΔR^2^ = 1.25%) and drinks per week in FT12 (ΔR^2^ = 1.17%), and PRS for alcohol problems (COGA = 1.18%; FT12 = 0.40%).

**Figure 1:**
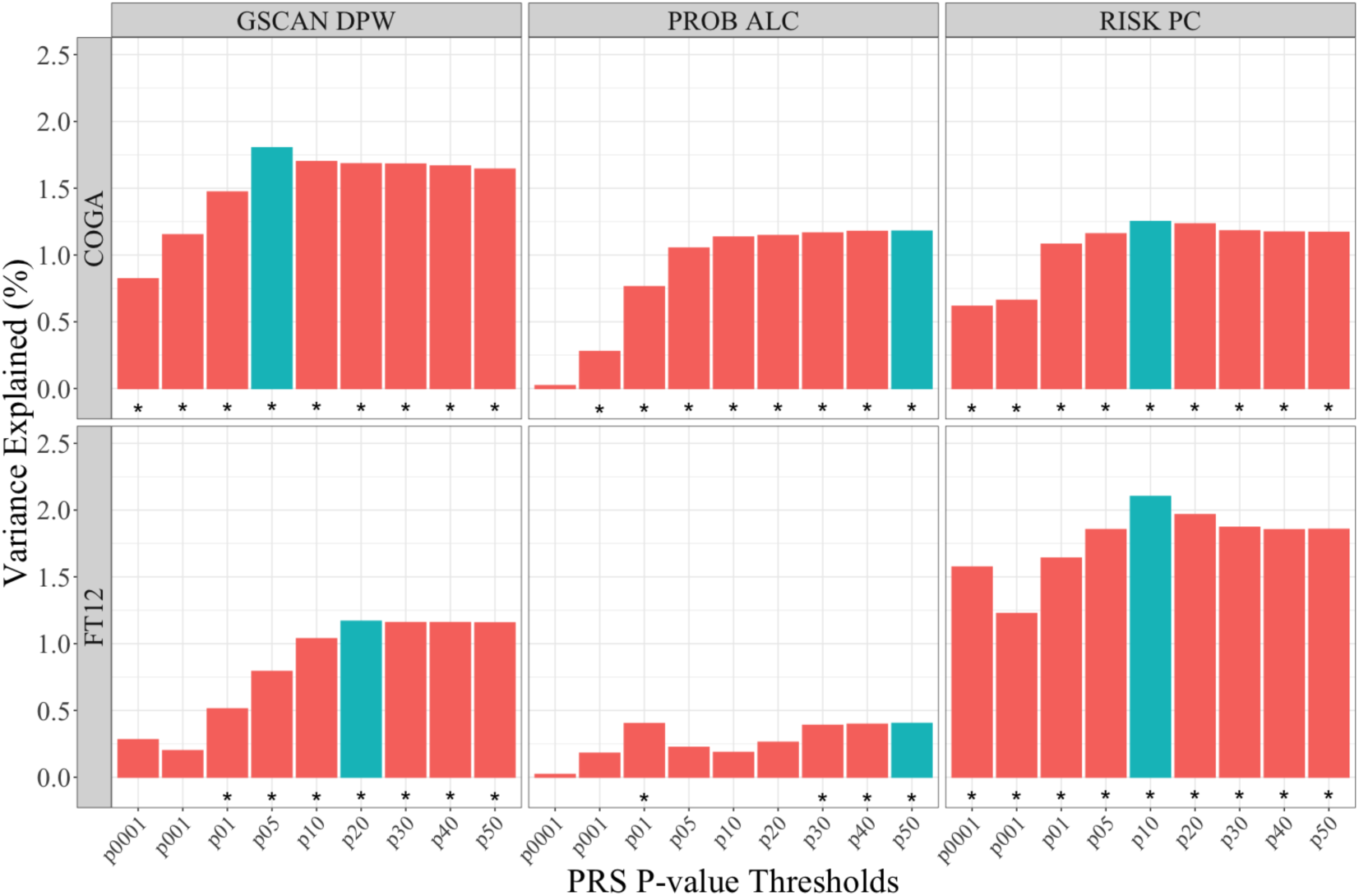
Predictive Power of PRS across Samples. Predictive power of PRS measured using pseudo-R^2^ for linear mixed effects models ^28^. Bars represent change in variance explained over models with age, sex, and first 10 ancestral principal components, genotyping array, and data collection site (only COGA for the latter two). Most predictive score outlined in blue. * p < .05, corrected for FDR of 5%

### Increase in Risk across the Polygenic Continuum

In order to estimate whether individuals at the extreme end of the PRS distribution were at elevated risk of AUD, we compared risk of AUD between those above and below a given threshold in the distribution. First, we determined whether each of these PRS contributed to AUD symptoms in a model containing all three, jointly. Figure 2 presents the parameter estimates for the independent and conditional effect of each PRS in both COGA and FT12. In the conditional model for COGA, each of the PRSs remains significantly associated with AUD symptoms, though the associations are attenuated (conditional model ΔR^2^ = 2.65%). In FT12, only the PRS for RISK PC remains significant in the joint model (conditional model ΔR^2^ = 2.45%). We averaged the three PRS into one composite PRS score of genetic risk in COGA and used the RISK PC PRS in FT12 to carry forward in the following analyses.

**Figure 2:**
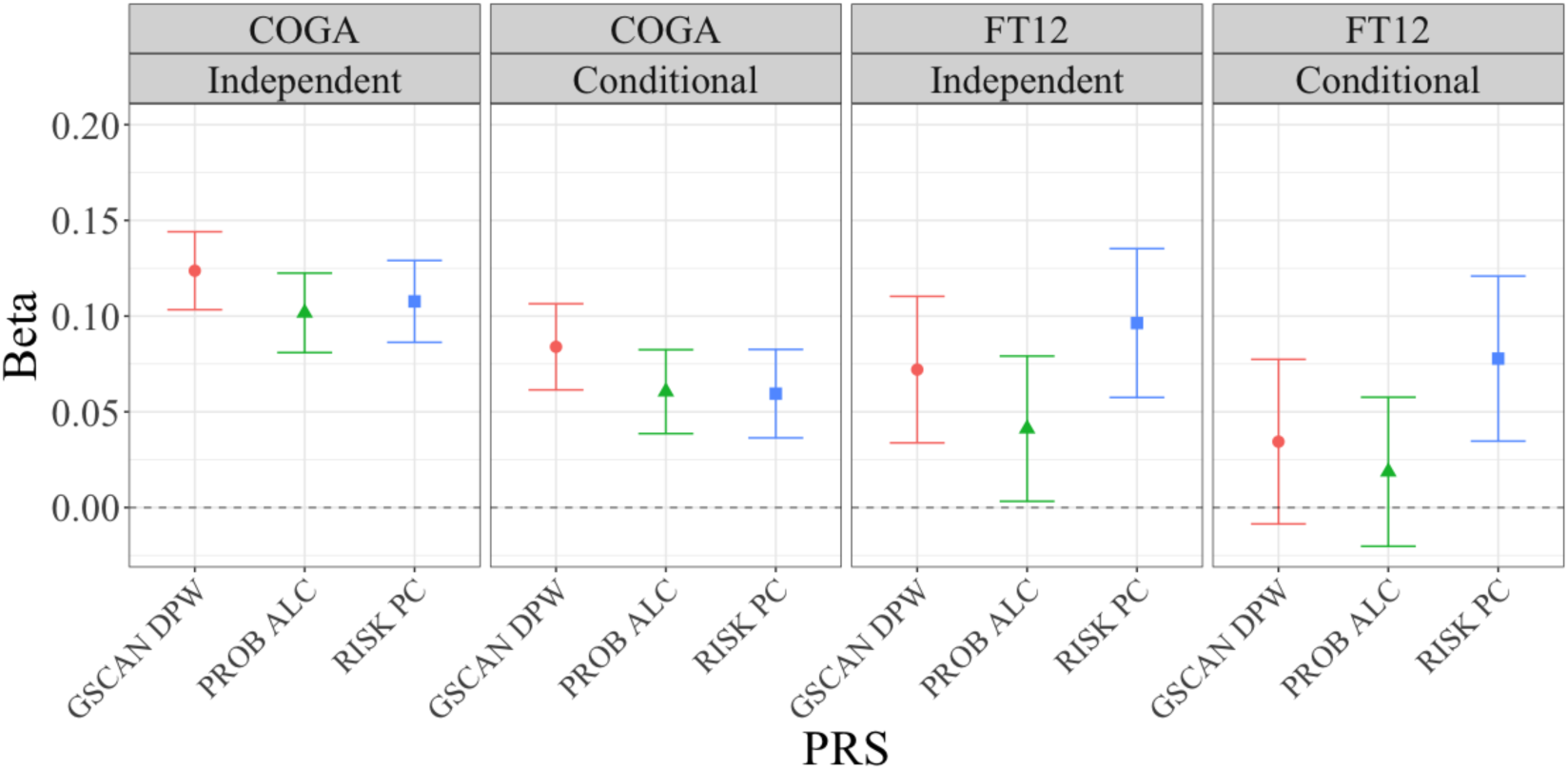
Parameter Estimates for PRS in Independent and Joint Models. Parameter estimates from linear mixed models for AUD symptoms regressed on GSCAN DPW, PROB ALC, and RISK PC PRS in COGA and FT12. Independent = model with only corresponding PRS. Conditional = model with all PRS included. Adjusted for age, sex, first 10 ancestral principal components, genotyping array, and data collection site (only COGA for the latter two).

Next, we divided these PRSs at the 80^th^, 90^th^, and 95^th^ percentile in each sample and estimated the odds ratio (OR) for AUD in the top portion of the distribution relative to the bottom portion of the distribution (e.g. splitting at the 80^th^ percentile compares the top 20% to the bottom 80%). Table 2 provides the estimates for all of those models. Across each threshold for AUD severity in COGA, we observed a similar pattern where restricting to the more extreme end of the polygenic distribution resulted in greater odds of meeting criteria for AUD. For example, there was increasing risk for a severe AUD when dividing 80^th^ percentile (OR = 1.948; 95% CI = 1.665, 2.278), 90^th^ percentile (OR = 2.027; 95% CI = 1.655, 2.482), and 95^th^ percentile (OR = 2.126; 95% CI = 1.617, 2.796). In FT12, there was a similar pattern for mild and severe AUD, but not moderate AUD. However, given the small number of cases in the extreme end for severe AUD, these estimates should be interpreted cautiously. Finally, we assessed the sensitivity/specificity of these PRS by calculating the AUC. AUC from the full model (including both continuous PRS and covariates) for each level of AUD severity ranged from 0.67 to 0.74 in COGA and from 0.65 to 0.75 in FT12. Comparing the AUC for the models with and without PRSs, including the PRS only increased the AUC slightly (see supplemental information).

**Table 2:** Odds Ratios for Those at Extreme End of the PRS Continuum.

| <u>Sample</u> | <u>Phenotype</u> | <u>Prevalence</u> | <u>Split</u> | <u>N Cases</u> | <u>OR</u> | <u>95 % CI Low</u> | <u>95 % CI High</u> |
| --- | --- | --- | --- | --- | --- | --- | --- |
| COGA | Mild AUD | 57.06% | 80% | 999 | 1.94 | 1.68 | 2.24 |
|  | Mild AUD | 57.06% | 90% | 522 | 1.97 | 1.62 | 2.39 |
|  | Mild AUD | 57.06% | 95% | 276 | 2.23 | 1.69 | 2.95 |
| COGA | Moderate AUD | 37.44% | 80% | 743 | 1.97 | 1.71 | 2.27 |
|  | Moderate AUD | 37.44% | 90% | 401 | 2.04 | 1.69 | 2.46 |
|  | Moderate AUD | 37.44% | 95% | 218 | 2.25 | 1.73 | 2.92 |
| COGA | Severe AUD | 25.89% | 80% | 547 | 1.95 | 1.67 | 2.28 |
|  | Severe AUD | 25.89% | 90% | 300 | 2.03 | 1.66 | 2.48 |
|  | Severe AUD | 25.89% | 95% | 165 | 2.13 | 1.62 | 2.80 |
| FT12 | Mild AUD | 41.98% | 80% | 122 | 1.77 | 1.22 | 2.56 |
|  | Mild AUD | 41.98% | 90% | 68 | 2.27 | 1.39 | 3.72 |
|  | Mild AUD | 41.98% | 95% | 36 | 2.39 | 1.21 | 4.72 |
| FT12 | Moderate AUD | 13.91% | 80% | 45 | 1.80 | 1.09 | 2.98 |
|  | Moderate AUD | 13.91% | 90% | 22 | 1.52 | 0.79 | 2.92 |
|  | Moderate AUD | 13.91% | 95% | 12 | 1.59 | 0.66 | 3.80 |
| FT12 | Severe AUD | 3.79% | 80% | 15 | 2.16 | 1.06 | 4.40 |
|  | Severe AUD | 3.79% | 90% | 9 | 2.45 | 1.05 | 5.71 |
|  | Severe AUD | 3.79% | 95% | 6 | 3.24 | 1.11 | 9.53 |

All models control for sex, age at last interview, and first 10 principal components. Models for COGA also included data collection site and genotyping array. N Cases = number of individuals who meet criteria for a given level of AUD and are in the top portion of the split.

### Examining the Substance Use Phenome of the Extreme End of the Polygenic Risk Continuum

We compared the likelihood of substance-related outcomes in individuals in the top 5% of each of the PRS in COGA and FT12 (adjusted for covariates). Figure 3 presents the mean lifetime symptoms endorsed for a variety of substance use disorders (alcohol, cannabis, cocaine, nicotine, and opioid) for individuals in the top 5% for each PRS relative to the bottom 95% of each PRS. In COGA, individuals in the top 5% of the PROB ALC, RISK PC, and/or GSCAN DPW PRS had significantly higher levels of alcohol (0.25 – 0.33 SD) and nicotine symptoms (0.13 – 0.18 SD) than those in the bottom 95% of the PRS distribution. Those in the top 5% of the RISK PC PRS also endorsed a higher number of cannabis use disorder symptoms (0.14 SD). In FT12, those in the top 5% did not differ significantly for AUD or FTND symptoms.

**Figure 3:**
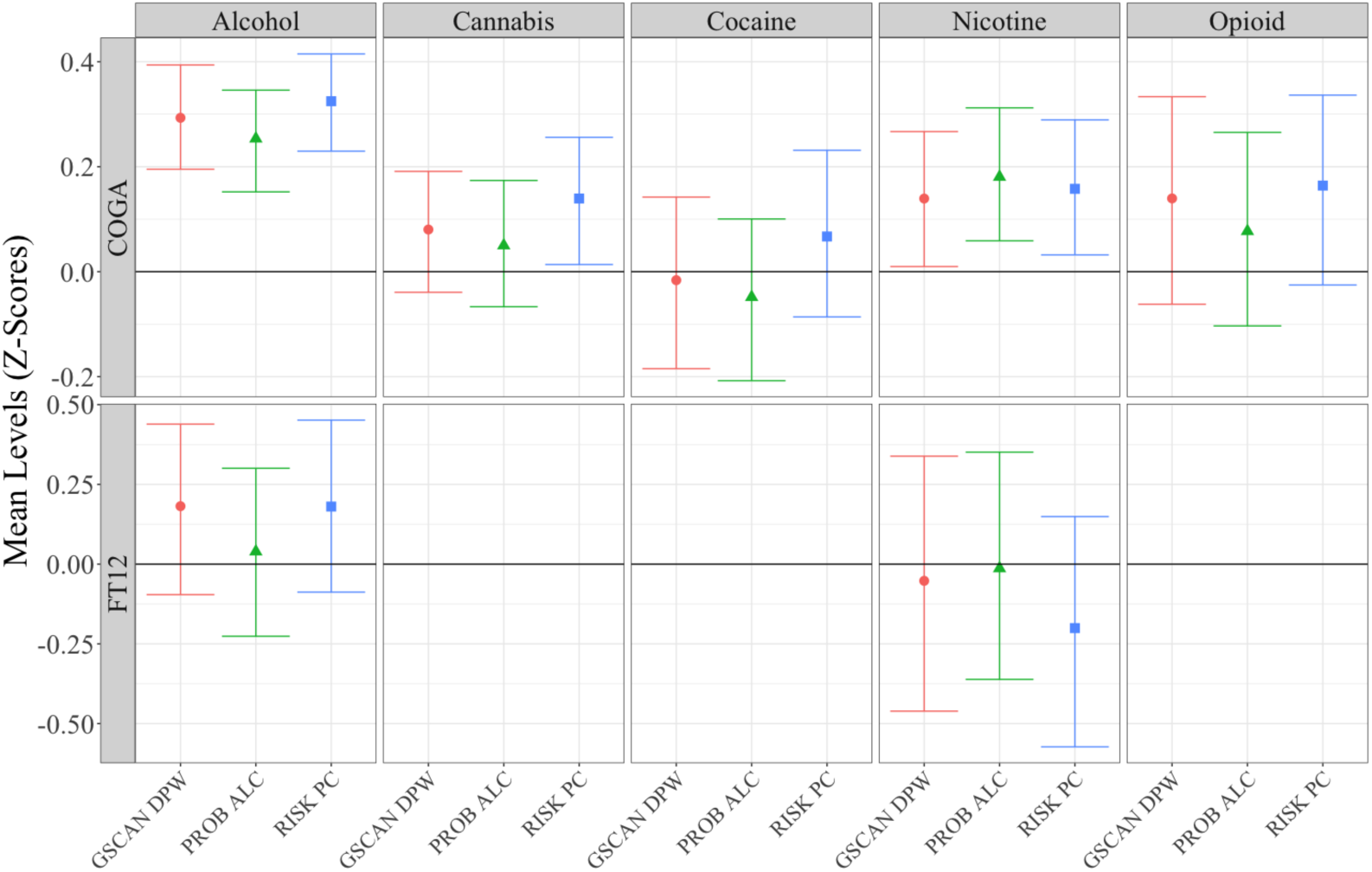
Top 5% of PRS Continuum. Mean levels of SUD symptoms for alcohol, cannabis, cocaine, nicotine, and opioid use disorders for top 5% of each PRS compared to the bottom 50%. Black bar represents mean of bottom 50%. 95% confidence intervals estimated using 1000 bootstrap resampling.

Overall, individuals in the top 5% of any PRS report greater levels of any substance, though being in the top 5% of the RISK PC PRS is associated with the most other substances. These PRS are modestly correlated with one another in both COGA (*r*_*RISK PC* PROB ALC*_ = .30; *r*_*GSCAN DPW*RISK PC*_ = .37; *r*_*GSCAN DPW* PROB*_ *ALC* = .32) and FT12 (*rRISK PC*PROB ALC* = .20; *rGSCAN DPW*RISK PC* = .45; *rGSCAN DPW* PROB ALC* = .27). These PRS each seem to capture unique information related to the genetics of substance use problems (and other risky behaviors).

## Discussion

Researchers have begun to evaluate the clinical utility of PRS for a variety of medical phenotypes 11, 12. In this analysis, we examined the current utility of several PRSs for identifying those at risk for a variety of SUDs, with a focus on AUD in both a clinically ascertained and population based sample. We were interested in 1) which scores based on available GWASs provide the best prediction for alcohol use disorder and whether these scores predicted unique variance in AUD in a joint model; 2) what the risk of AUD was for those at the upper end of the risk continuum compared to the bottom; and 3) the levels of substance use disorder symptoms for individuals at the top 5% of the polygenic score continuum compared to remaining 95%.

In terms of which polygenic scores were the most predictive, we considered three scores: one based on problematic alcohol use (PROB ALC), one based on alcohol consumption (GSCAN DPW), and one based on general risky behaviors (RISK PC), as twin and family studies have shown alcohol and other risk behaviors to be genetically correlated traits ^6, 14-18^. In the population sample (FT12), the RISK PC PRS was the most predictive. These results support the idea that focusing on the shared genetic etiology towards risk taking, sometimes referred to as externalizing ^16-18^, may prove useful for identifying those at risk for SUD ^30^. In our more clinically based (COGA) sample, the PRS for alcohol consumption (GSCAN DPW) explained the largest portion of the variance. When we included all of the PRS in one model, all three PRS were associated with AUD symptoms in COGA while only the RISK PC PRS was associated with AUD symptoms in FT12. We ran a series of sensitivity analyses to test whether differences across the samples reflected age differences rather than differences in ascertainment. Restricting COGA to participants under 30 did not fundamentally change the results.

When we divided the PRS at different thresholds, the odds of having an AUD steadily increased from the 80% threshold to the 95% threshold in both COGA and FT12. However, even though the point estimates steadily increased, the confidence intervals around these estimates were relatively large and they did not differ significantly. Additionally, there were only a small number of individuals in the severe category in FT12 and we urge caution in interpreting these estimates. Supplemental analyses evaluating the AUC for each continuous PRS in a joint model revealed the combined effect of all three PRSs only marginally improved the AUC over models with just covariates. Finally, the top 5% of the continuum for each PRS reported elevated rates of other SUD symptoms (cannabis, cocaine, and nicotine use disorders) compared to the bottom 95%. The RISK PC PRS was the most predictive of these higher mean levels of SUD symptoms, suggesting that risk for externalizing may be particularly useful in identifying individuals at risk for SUDs.

These initial findings suggest the use of genetic data may eventually have utility in a clinical setting for SUDs, but not in their current form. Being able to identify those at heightened risk for SUDs may allow for more targeted early intervention and prevention. Before clinical utility is possible, larger discovery GWAS across substance use phenotypes with PRS that explain greater portions of the variance will be necessary. As GWAS sample sizes for SUDs increase, we will likely see improved prediction ^31^. Additionally, using multivariate techniques to model the shared genetic architecture across existing SUD GWAS to include both aspects of externalizing and internalizing (e.g. depression, anxiety) may also improve prediction ^32, 33^. Inclusion of genetic data in a clinical setting will also require that psychiatrists and clinicians receive greater training in genetics and/or that they partner with genetic counselors, so they are both better able to understand what increased genetic risk means and be able convey that information accurately to their patients ^34, 35^. In addition to clinical utility, we must ensure that regulations and protections surrounding the use of genetic information in clinical settings can adequately protect the rights of individuals who are identified to be “at risk.”

This research has several important limitations. First, all analyses were limited to individuals of European ancestry because the discovery GWASs available were conducted in individuals of primarily European ancestry. It will be important to ascertain sizable samples of subjects with non-European ancestries to properly estimate the predictive utility of PRS in non-European samples. This is especially important for racial-ethnic minorities so that health disparities are not further perpetuated ^36^. Second, our use of lifetime diagnoses may obscure the impact of changing genetic influences on the development of AUD across the life course ^37, 38^. Future work should draw on longitudinal data to examine the ways in which the predictive utility of PRS changes with the age of the target sample. Finally, these analyses examined the marginal influence of PRS, independent of environment. Processes of gene-environment interaction (GxE) are well documented in alcohol misuse using twin and family data ^39-42^. Incorporating environmental information along with PRS in a methodologically rigorous manner will be an important next step in developing clinically predictive algorithms.

Genome-wide polygenic scores are beginning to have utility in identifying individuals at risk of certain diseases, especially those related to well defined physical health conditions, such as cardiovascular disease ^11, 12^. We examined the current state of PRS for predicting substance use, with a focus on AUD. Each of the PRS analyzed here predicted AUD. However, the overall maximum variance explained was still small (∼2%). Individuals at the top of the PRS continuum had elevated rates of multiple substance use problems, but these differences across the PRS continuum are unlikely to be of broad clinical use in their current state. As GWAS discovery samples become larger and we are better able to model the complex relationship between genotype and phenotype, polygenic scores may eventually be useful in a clinical setting.

## Acknowledgments

Research reported in this publication was supported by the National Institute on Alcohol Abuse and Alcoholism of the National Institutes of Health under award numbers R01AA015416 (DMD), K02AA018755 (DMD), K02DA32573 (AA), and F32AA027435 (ECJ); the Academy of Finland (grants 100499, 205585, 118555, 141054, 265240, 308248, 308698 and 312073); and the Scientific and Technological Research Council of Turkey (TÜBİTAK) under award number 114C117 (FA). The content is solely the responsibility of the authors and does not necessarily represent the official views of the National Institutes of Health, the Academy of Finland, or the Scientific and Technological Research Council of Turkey. This research also used summary data from the Psychiatric Genomics Consortium (PGC) Substance Use Disorders (SUD) working group. The PGC-SUD is supported by funds from NIDA and NIMH to MH109532 and, previously, had analyst support from NIAAA to U01AA008401 (COGA). PGC-SUD gratefully acknowledges its contributing studies and the participants in those studies, without whom this effort would not be possible.

## COGA

The Collaborative Study on the Genetics of Alcoholism (COGA), Principal Investigators B. Porjesz, V. Hesselbrock, H. Edenberg, L. Bierut, includes eleven different centers: University of Connecticut (V. Hesselbrock); Indiana University (H.J. Edenberg, J. Nurnberger Jr., T. Foroud; Y. Liu); University of Iowa (S. Kuperman, J. Kramer); SUNY Downstate (B. Porjesz); Washington University in St. Louis (L. Bierut, J. Rice, K. Bucholz, A. Agrawal); University of California at San Diego (M. Schuckit); Rutgers University (J. Tischfield, A. Brooks); Department of Biomedical and Health Informatics, The Children’s Hospital of Philadelphia; Department of Genetics, Perelman School of Medicine, University of Pennsylvania, Philadelphia PA (L. Almasy), Virginia Commonwealth University (D. Dick), Icahn School of Medicine at Mount Sinai (A. Goate), and Howard University (R. Taylor). Other COGA collaborators include: L. Bauer (University of Connecticut); J. McClintick, L. Wetherill, X. Xuei, D. Lai, S. O’Connor, M. Plawecki, S. Lourens (Indiana University); G. Chan (University of Iowa; University of Connecticut); J. Meyers, D. Chorlian, C. Kamarajan, A. Pandey, J. Zhang (SUNY Downstate); J.-C. Wang, M. Kapoor, S. Bertelsen (Icahn School of Medicine at Mount Sinai); A. Anokhin, V. McCutcheon, S. Saccone (Washington University); J. Salvatore, F. Aliev, B. Cho (Virginia Commonwealth University); and Mark Kos (University of Texas Rio Grande Valley). A. Parsian and H. Chen are the NIAAA Staff Collaborators.

We continue to be inspired by our memories of Henri Begleiter and Theodore Reich, founding PI and Co-PI of COGA, and also owe a debt of gratitude to other past organizers of COGA, including Ting-Kai Li, P. Michael Conneally, Raymond Crowe, and Wendy Reich, for their critical contributions. This national collaborative study is supported by NIH Grant U10AA008401 from the National Institute on Alcohol Abuse and Alcoholism (NIAAA) and the National Institute on Drug Abuse (NIDA).

## References

1. Gakidou E, Afshin A, Abajobir AA, Abate KH, Abbafati C, Abbas KM et al. Global, regional, and national comparative risk assessment of 84 behavioural, environmental and occupational, and metabolic risks or clusters of risks, 1990–2016: a systematic analysis for the Global Burden of Disease Study 2016. The Lancet 2017; 390(10100): 1345–1422.

2. The USBoDC. The state of us health, 1990-2016: Burden of diseases, injuries, and risk factors among us states. JAMA 2018; 319(14): 1444–1472.

3. World Health Organization. Global status report on alcohol and health - 2018. Geneva, Switzerland 2018.

4. Verhulst B, Neale MC, Kendler KS. The heritability of alcohol use disorders: a meta-analysis of twin and adoption studies. Psychological Medicine 2015; 45(5): 1061–1072.

5. Walters RK, Polimanti R, Johnson EC, McClintick JN, Adams MJ, Adkins AE et al. Transancestral GWAS of alcohol dependence reveals common genetic underpinnings with psychiatric disorders. Nature Neuroscience 2018; 21(12): 1656–1669.

6. Sanchez-Roige S, Palmer AA, Fontanillas P, Elson SL, Adams MJ, Howard DM et al. Genome-Wide Association Study Meta-Analysis of the Alcohol Use Disorders Identification Test (AUDIT) in Two Population-Based Cohorts. American Journal of Psychiatry 2018: appi.ajp.2018.18040369.

7. Kranzler HR, Zhou H, Kember RL, Vickers Smith R, Justice AC, Damrauer S et al. Genome-wide association study of alcohol consumption and use disorder in 274,424 individuals from multiple populations. Nat Commun 2019; 10(1): 1499.

8. Liu M, Jiang Y, Wedow R, Li Y, Brazel DM, Chen F et al. Association studies of up to 1.2 million individuals yield new insights into the genetic etiology of tobacco and alcohol use. Nature Genetics 2019.

9. Gelernter J, Sun N, Polimanti R, Pietrzak R, Levey DF, Lu Q et al. Genomewide Association Study of Maximum Habitual Alcohol Intake in >140,000 US European- and African-American Veterans Yields Novel Risk Loci. Biological Psychiatry 2019.

10. Pasman JA, Verweij KJH, Gerring Z, Stringer S, Sanchez-Roige S, Treur JL et al. GWAS of lifetime cannabis use reveals new risk loci, genetic overlap with psychiatric traits, and a causal influence of schizophrenia. Nature Neuroscience 2018; 21(9): 1161–1170.

11. Khera AV, Chaffin M, Aragam KG, Haas ME, Roselli C, Choi SH et al. Genome-wide polygenic scores for common diseases identify individuals with risk equivalent to monogenic mutations. Nature Genetics 2018; 50(9): 1219–1224.

12. Khera AV, Chaffin M, Wade KH, Zahid S, Brancale J, Xia R et al. Polygenic Prediction of Weight and Obesity Trajectories from Birth to Adulthood. Cell 2019; 177(3): 587–596 e589.

13. Karlsson Linnér R, Biroli P, Kong E, Meddens SFW, Wedow R, Fontana MA et al. Genome-wide association analyses of risk tolerance and risky behaviors in over 1 million individuals identify hundreds of loci and shared genetic influences. Nature Genetics 2019; 51(2): 245–257.

14. Dick DM, Meyers JL, Rose RJ, Kaprio J, Kendler KS. Measures of Current Alcohol Consumption and Problems: Two Independent Twin Studies Suggest a Complex Genetic Architecture. Alcoholism: Clinical and Experimental Research 2011; 35(12): 2152–2161.

15. Kendler KS, Myers J, Dick D, Prescott CA. The Relationship Between Genetic Influences on Alcohol Dependence and on Patterns of Alcohol Consumption. Alcoholism: Clinical and Experimental Research 2010; 34(6): 1058–1065.

16. Kendler KS, Myers J. The boundaries of the internalizing and externalizing genetic spectra in men and women. Psychological Medicine 2013; 44(3): 647–655.

17. Krueger RF, Hicks BM, Patrick CJ, Carlson SR, Iacono WG, McGue M. Etiologic connections among substance dependence, antisocial behavior and personality: Modeling the externalizing spectrum. Journal of Abnormal Psychology 2002; 111(3): 411–424.

18. Kendler KS, Prescott CA, Myers J, Neale MC. The structure of genetic and environmental risk factors for common psychiatric and substance use disorders in men and women. Archives of General Psychiatry 2003; 60(9): 929–937.

19. Bucholz KK, Cadoret R, Cloninger CR, Dinwiddie SH, Hesselbrock VM, Nurnberger JI et al. A new, semi-structured psychiatric interview for use in genetic linkage studies: a report on the reliability of the SSAGA. Journal of Studies on Alcohol 1994; 55(2): 149–158.

20. Kaprio J. The Finnish Twin Cohort Study: an update. Twin Res Hum Genet 2013; 16(1): 157–162.

21. Bucholz KK, McCutcheon VV, Agrawal A, Dick DM, Hesselbrock VM, Kramer J et al. Comparison of Parent, Peer, Psychiatric, and Cannabis Use Influences Across Stages of Offspring Alcohol Involvement: Evidence from the COGA Prospective Study. Alcoholism: Clinical and Experimental Research 2017; 41(2): 359–368.

22. Edenberg HJ. The collaborative study on the genetics of alcoholism: an update. Alcohol research & health: the journal of the National Institute on Alcohol Abuse and Alcoholism 2002; 26(3): 214–218.

23. Begleiter H, Reich T, Hesselbrock V, Porjesz B, Li T-K, Schuckit MA et al. The collaborative study on the genetics of alcoholism. Alcohol Health and Research World 1995; 19: 228–228.

24. Martin AR, Gignoux CR, Walters RK, Wojcik GL, Neale BM, Gravel S et al. Human Demographic History Impacts Genetic Risk Prediction across Diverse Populations. Am J Hum Genet 2017; 100(4): 635–649.

25. American Psychiatric Association. Diagnostic and statistical manual of mental disorders (DSM-5®). American Psychiatric Pub 2013.

26. International Schizophrenia Consortium. Common polygenic variation contributes to risk of schizophrenia and bipolar disorder. Nature 2009; 460(7256): 748–752.

27. Purcell S, Neale B, Todd-Brown K, Thomas L, Ferreira MA, Bender D et al. PLINK: a tool set for whole-genome association and population-based linkage analyses. Am J Hum Genet 2007; 81(3): 559–575.

28. Nakagawa S, Schielzeth H, O’Hara RB. A general and simple method for obtaining R-squared from generalized linear mixed-effects models. Methods in Ecology and Evolution 2013; 4(2): 133–142.

29. Wald NJ, Old R. The illusion of polygenic disease risk prediction. Genetics in Medicine 2019.

30. Dick DM, Aliev F, Wang JC, et al. Using dimensional models of externalizing psychopathology to aid in gene identification. Archives of General Psychiatry 2008; 65(3): 310–318.

31. Dudbridge F. Power and Predictive Accuracy of Polygenic Risk Scores. PLOS Genetics 2013; 9(3): e1003348.

32. Dick DM, Koellinger P, Harden KP, Tucker-Drob EM, Waldman I, Karlsson Linnér R et al. Using the Genetic Architecture of Externalizing Disorders to Aid in Gene Identification. Annual Meeting of the Behavior Genetics Association: Boston, MA., 2018.

33. Grotzinger AD, Rhemtulla M, de Vlaming R, Ritchie SJ, Mallard TT, Hill WD et al. Genomic structural equation modelling provides insights into the multivariate genetic architecture of complex traits. Nature Human Behaviour 2019.

34. Besterman AD, Moreno-De-Luca D, Nurnberger JI, Jr. 21st-Century Genetics in Psychiatric Residency Training: How Do We Get There? JAMA Psychiatry 2019.

35. Nurnberger JI, Jr., Austin J, Berrettini WH, Besterman AD, DeLisi LE, Grice DE et al. What Should a Psychiatrist Know About Genetics? Review and Recommendations From the Residency Education Committee of the International Society of Psychiatric Genetics. The Journal of clinical psychiatry 2018; 80(1).

36. Martin AR, Kanai M, Kamatani Y, Okada Y, Neale BM, Daly MJ. Clinical use of current polygenic risk scores may exacerbate health disparities. Nature Genetics 2019; 51(4): 584–591.

37. Kendler KS, Gardner C, Dick D. Predicting alcohol consumption in adolescence from alcohol-specific and general externalizing genetic risk factors, key environmental exposures and their interaction. Psychological Medicine 2011; 41(07): 1507--1516.

38. Meyers JL, Salvatore JE, Vuoksimaa E, Korhonen T, Pulkkinen L, Rose RJ et al. Genetic Influences on Alcohol Use Behaviors Have Diverging Developmental Trajectories: A Prospective Study Among Male and Female Twins. Alcoholism: Clinical and Experimental Research 2014; 38(11): 2869–2877.

39. Barr PB, Kuo SI, Aliev F, Latvala A, Viken R, Rose RJ et al. Polygenic Risk for Alcohol Misuse is Moderated by Romantic Partnerships. Addiction 2019.

40. Cooke ME, Meyers JL, Latvala A, Korhonen T, Rose RJ, Kaprio J et al. Gene-Environment Interaction Effects of Peer Deviance, Parental Knowledge and Stressful Life Events on Adolescent Alcohol Use. Twin Research and Human Genetics 2015; 18(5): 507–517.

41. Dick DM, Bernard M, Aliev F, Viken R, Pulkkinen L, Kaprio J et al. The role of socioregional factors in moderating genetic influences on early adolescent behavior problems and alcohol use. Alcoholism: Clinical and Experimental Research 2009; 33(10): 1739–1748.

42. Dick DM, Viken R, Purcell S, Kaprio J, Pulkkinen L, Rose RJ. Parental monitoring moderates the importance of genetic and environmental influences on adolescent smoking. Journal of Abnormal Psychology 2007; 116(1): 213–218.

