## Supplemental information for "Polygenic Prediction of Substance Use Disorders in Clinical and Population Samples"

### 1. Genotyping and Quality Control

#### *COGA*

Genotyping of the COGA samples was conducted across different phases of data collection. European ancestry (EA) samples were genotyped at multiple sites, including: (1) Center for Inherited Disease Research using the Illumina HumanHap1M array [1]; (2) Genome Technology Access Center at Washington University School of Medicine using the Illumina OmniExpress [2]; and (3) Rutgers University using the Affymetrix Smokescreen array [3]. All A/T and C/G SNPs were removed and a common set of ~47,000 SNPs were used to assess duplicate samples and revise the reported pedigree structure. Family structures were altered as needed, and SNP genotypes were tested for Mendelian inconsistencies [4] with the revised family structure. Genotype inconsistencies were set to missing. Data were imputed to 1000 Genomes (Phase 3, version 5) using SHAPEIT [5] and then Minimac3 [6]. Samples were imputed separately due to different variant contents on each array. In addition, the two datasets genotyped on the Smokescreen genotyping array were also imputed separately, due to different processing pipelines used by the genotyping laboratory. Prior to imputation, variants with missing rates > 5%, MAF < 3% and HWE p values < 0.0001 were excluded, resulting in a total of 26,124,746 variants. After QC, there were 3,440,839 in common. Of these variants, 367,710 autosomal SNPs were extracted for creating PRS after LD clumping. Principal components were computed from GWAS data using Eigenstrat [7] and 1000 Genome reference panel. Individual ancestry was assigned using the YRI, CEU, JPT and CHB populations to set reference points.

#### *FinnTwin12*

Genotyping was conducted using the Human670-QuadCustom Illumina BeadChip at the Wellcome Trust Sanger Institute [8]. Quality control steps included removing SNPs with minor allele frequency (MAF) <1%, genotyping success rate <95%, or Hardy–Weinberg equilibrium $p < 1 \times 10^{-6}$, and removing individuals with genotyping success rate <95%, a mismatch between phenotypic and genotypic gender, excess relatedness (outside of known families), and heterozygosity outliers. Genotypes were imputed to the 1,000 Genomes Phase 3 reference panel [9] reference panel using ShapeIT [10] for phasing and IMPUTE2 [11] for imputation, resulting in 13,688,418 autosomal SNPs for analyses. There were 3,336,209 autosomal SNPs in common after QC and 346,604 independent SNPs for creating scores after LD clumping.

#### 2. Problematic Alcohol use Meta Analysis

In order to maximize the predictive power for polygenic scores (PRS) for problematic alcohol use, we meta-analyzed summary statistics from two recent large scale GWAS of problematic alcohol use. The first GWAS was conducted using the problem subscale of the Alcohol Use Disorder Identification Test (AUDIT-P) on ~120k individuals in the UK Biobank. The AUDIT-P is made up of items 4-10 (1-3 reflect consumptions, or AUDIT-C) of the full AUDIT. Scores on individual items range from 0 to 4, resulting in an overall range of 0 to 28 for the AUDIT-P. Items include:

1. How often during the last year have you found that you were unable to stop drinking once you had started?
2. How often during the last year have you failed to do what was normally expected of you because of your drinking
3. How often during the last year have you needed a first drink in the morning to get yourself going after a heavy drinking session?
4. How often during the last year have you had a feeling of guilt or remorse after drinking?
5. How often during the last year have you been unable to remember what happened the night before because you had been drinking?
6. Have you or someone else been injured as the result of your drinking?
7. Has a relative, friend, or a doctor or other health worker been concerned about your drinking or suggested you cut down?

The second GWAS was the Psychiatric Genetics Consortium's (PGC) GWAS of alcohol dependence (AD) in ~45K individuals (European ancestry results only). This GWAS focused on lifetime diagnosis (or meeting the criteria for lifetime diagnosis on a clinical interview) of AD. As both COGA and FT12 were included in the initial PGC GWAS, we recalculated summary statistics with each cohort removed. The resulting meta-analyses was a GWAS of problematic alcohol use in ~160K individuals.

We first estimate the SNP-based heritability ($h^2_{SNP}$) and genetic correlations (rg) between each of the GWAS (UKB AUDIT-P, PGC AD with COGA removed, and PGC AD with FT12 removed) using LD score regression [12]. Table S1 provides the estimates for $h^2_{SNP}$ and rg. Both of the PGC AD GWAS were sufficiently correlated with UKB AUDIT-P to justify meta-analysis.

To combine the results from these two sets of PGC summary statistics with the UKB AUDIT-P GWAS, we utilized a sample size based meta-analysis in METAL [13]. In addition to the quality control metrics in the original GWASs, we constrained to SNPS with a MAF > 0.01. Independent SNPs (r2 < 0.1 ) that met genome-wide significance (GWS) for each meta-analysis are presented in Table S2. For the NO COGA META, there were 13 significant SNPs, all on chromosome 4. The significant SNPs for the NO FT12 META were also on chromosome 4, however there was one additional independent SNP. In addition to these top SNPs, the Manhattan plots and QQ plots are available in Figures S1 and S2.

Follow up with the meta-analyzed results in LDSC revealed significant h2SNP in the both meta-analyses with COGA excluded (META NO COGA $h^2_{SNP}$ = 0.0565; SE = 0.0039; p = 1.46e-47) and with FT12 excluded (META NO FT12 $h^2_{SNP}$ = 0.058; SE = 0.0039; p = 5.02e-50). We see modest inflation in the test statistics (META NO COGA Mean $\chi^2$ = 1.1656; META NO FT12 Mean $\chi^2$ = 1.174). This genomic inflation appears to be the result of polygenic signal rather than population stratification, as the LDSC intercept is near one for each meta-analysis (META NO COGA Intercept = 1.0105, SE = 0.0065; META NO FT12 Intercept = 1.0108, SE = 0.0064). These GWAS meta-analysis results have a very high genetic correlation (rg = 0.9783, SE = 0.0017)

Table S1: LDSC Estimates for GWAS included in Meta-Analysis

| | | $h^2$ | | | rg (SE) | | |
|---|---|---|---|---|---|---|---|
| | **GWAS** | $h^2_{SNP}$ | **SE** | **p** | **1** | **2** | **3** |
| 1 | UKB AUDIT-P | 0.0573 | 0.0050 | 2.15e-05 | - | - | - |
| 2 | PGC (no COGA) | 0.0888 | 0.0209 | 7.17e-08 | 0.5869 (0.1229) | - | - |
| 3 | PGC (no FT12) | 0.0975 | 0.0181 | 2.10e-30 | 0.6349 (0.1137) | 0.9825 (0.0221) | - |

Table S2: Top SNPs from Meta-Analysis of PGC and UKB

| | | | | NO COGA META | | | NO FT12 META | | |
|---|---|---|---|---|---|---|---|---|---|
| **CHR** | **POS** | **RSID** | **Nearest Gene** | **Z** | **P** | **MAF** | **Z** | **P** | **MAF** |
| 4 | 100239319 | rs1229984 | ADH1B | -6.77 | 1.26E-11 | 0.03 | -7.13 | 9.78E-13 | 0.03 |
| 4 | 100244221 | rs3811802 | ADH1B | -8.02 | 1.02E-15 | 0.47 | -8.13 | 4.18E-16 | 0.47 |
| 4 | 100252560 | rs3114045 | ADH1C | - | - | - | -5.61 | 2.08E-08 | 0.13 |
| 4 | 100262242 | rs141973904 | ADH1C | -5.55 | 2.87E-08 | 0.02 | -6.40 | 1.52E-10 | 0.02 |
| 4 | 100282103 | rs4699743 | ADH1C | -5.97 | 2.30E-09 | 0.08 | -6.33 | 2.44E-10 | 0.08 |
| 4 | 103385336 | rs531685993 | AF213884.1 | 5.75 | 8.89E-09 | 0.09 | - | - | - |
| 4 | 99713350 | rs144198753 | BTF3P13 | -9.73 | 2.33E-22 | 0.01 | -9.73 | 2.33E-22 | 0.01 |
| 4 | 39411407 | rs62310819 | KLB | -5.47 | 4.50E-08 | 0.20 | -5.52 | 3.38E-08 | 0.20 |
| 4 | 39413780 | rs28712821 | KLB | 6.59 | 4.27E-11 | 0.39 | - | - | - |
| 4 | 39414993 | rs11940694 | KLB | - | - | - | -6.67 | 2.62E-11 | 0.39 |
| 4 | 99941138 | rs146788033 | METAP1 | 9.57 | 1.03E-21 | 0.02 | 9.71 | 2.77E-22 | 0.02 |
| 4 | 39393801 | rs6842066 | RNU6-887P | 6.00 | 2.01E-09 | 0.42 | - | - | - |
| 4 | 39400998 | rs13110790 | RNU6-887P | - | - | - | 6.74 | 1.58E-11 | 0.42 |
| 4 | 99630017 | rs193099203 | RP11-1299A16.1 | -8.08 | 6.61E-16 | 0.01 | -8.08 | 6.61E-16 | 0.01 |
| 4 | 100186847 | rs138423208 | RP11-696N14.1 | 6.70 | 2.16E-11 | 0.05 | 6.60 | 4.20E-11 | 0.05 |
| 4 | 103198082 | rs13135092 | SLC39A8 | 7.77 | 7.69E-15 | 0.09 | 7.92 | 2.39E-15 | 0.09 |
| 4 | 103283117 | rs34333163 | SLC39A8 | - | - | - | 6.06 | 1.33E-09 | 0.08 |

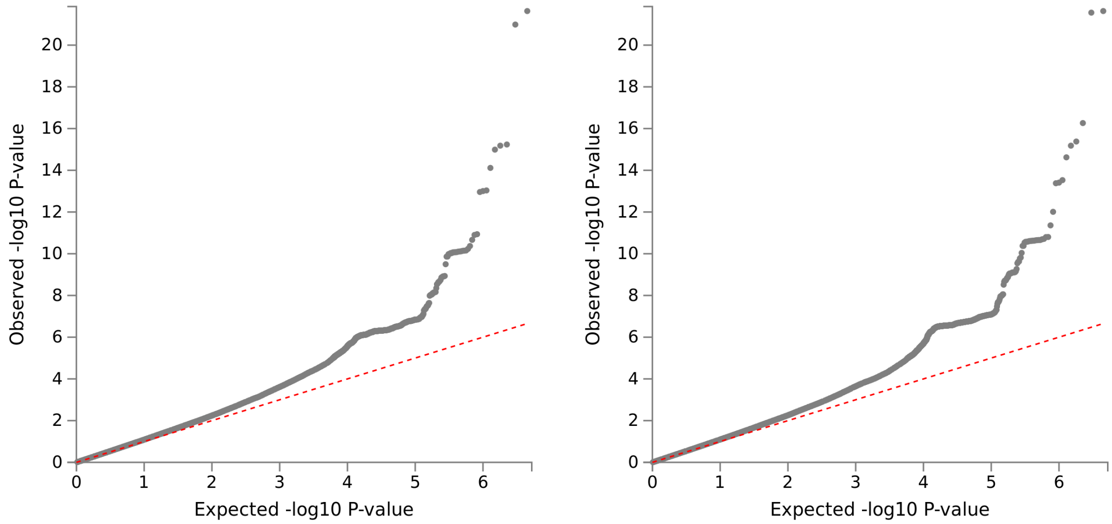


**Figure S1: QQ Plots for UK Biobank and PGC GWAS Meta Analyses**

QQ plots for observed versus expected p-values in meta analysis results with COGA (left) and FT12 (right) excluded.

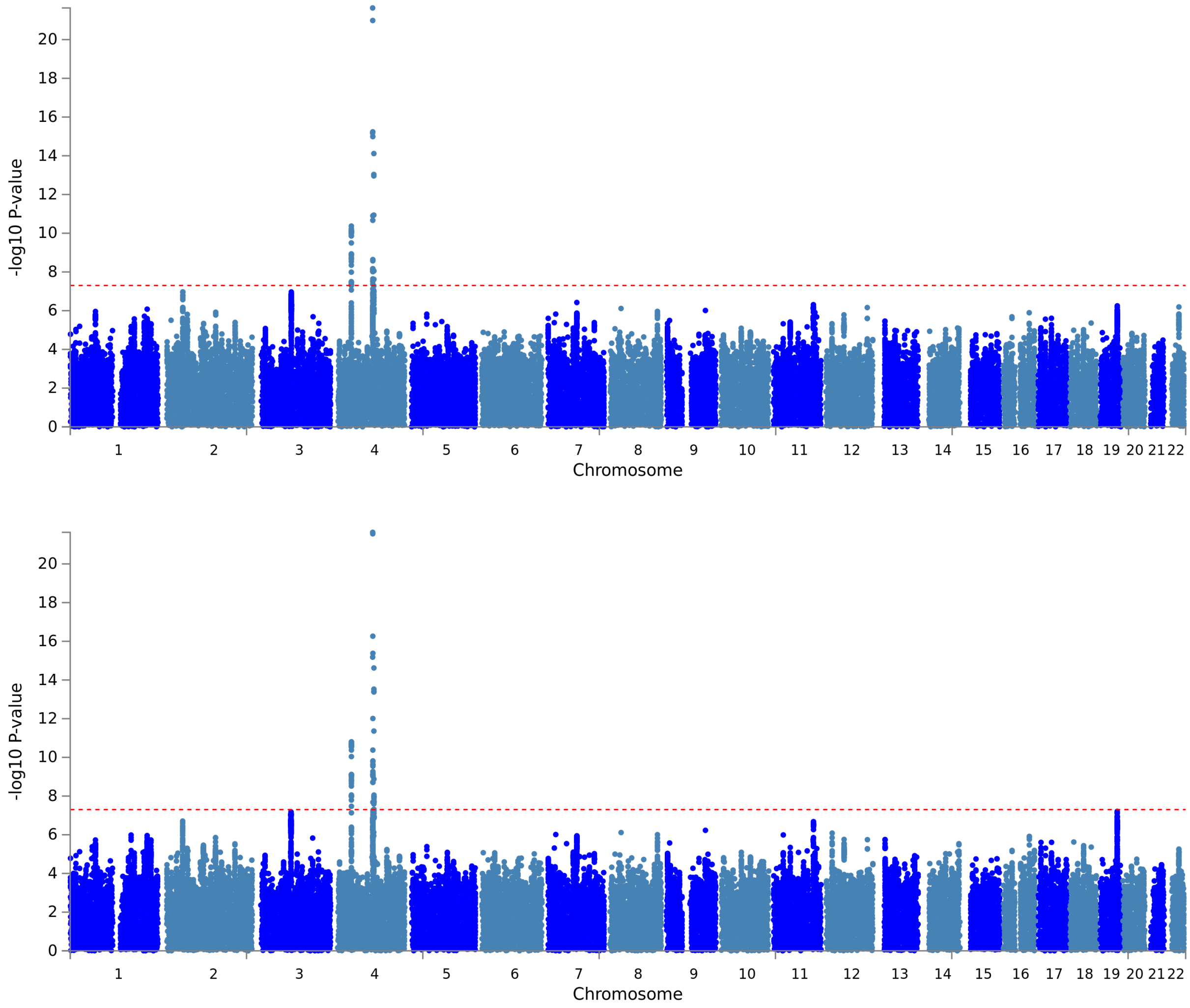


**Figure S2: Manhattan Plots for UK Biobank and PGC GWAS Meta Analyses**

Manhattan plots of –log$^{10}$ p-values for SNPS in meta-analysis with COGA (left) and FT12 (right) excluded.

### 3. Visualizing Increase in Risk across the Polygenic Continuum

We divided the most predictive PRS for each GWAS into septiles and plotted the prevalence of individuals meeting varying levels of AUD severity (mild, moderate, and severe) by PRS septile to determine if the relationship between PRS and AUD followed a consistently linear pattern. Results are shown in Figure 2. The linear increase in prevalence of AUD across the PRS risk continuum is evident in COGA for each of the PRS at each AUD severity threshold. As this sample is larger and contains many affected individuals with AUD, we are better able to identify associations between PRS and AUD. For FT12, there is a general increase in the prevalence of AUD across septiles of for the RISK PC and GSCAN DPW PRSs. However, for the PROB ALC PRS there does not appear to be an increase across septiles for moderate-to-severe AUD, reiterating the weaker predictive power of this PRS in FT12.

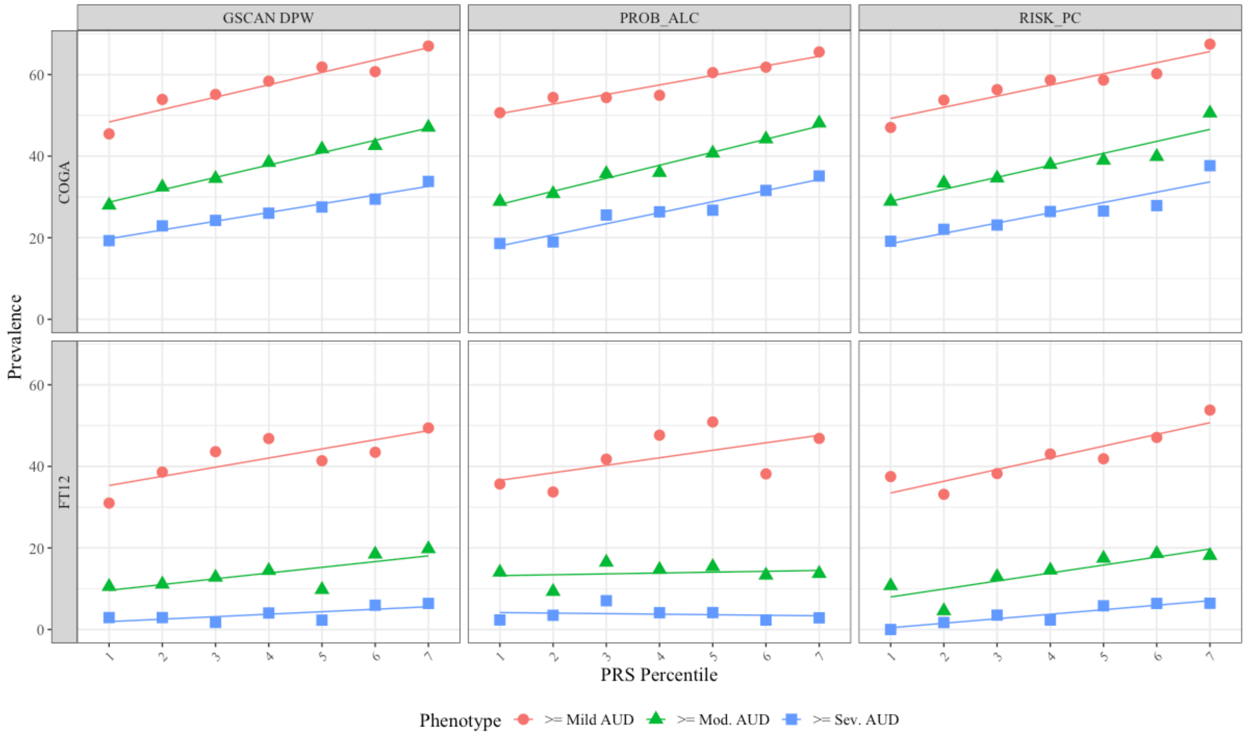


**Figure S3: Prevalence of AUD Across PRS Continuum**

Mean Prevalence of mild, moderate, and severe AUD in COGA and FT12 across deciles for GSCAN DPW, PROB ALC, and RISK PC PRS. Adjusted for age, sex, first 10 ancestral principal components, genotyping array, and data collection site (only COGA for the latter two).

### 4. Calculating AUC for PRS

While there is increasing focus on using the extreme ends of the polygenic risk continuum to as a potential way of identifying those at increased risk [14, 15], this method is not without its limitations. One critique, is that even though individuals at the top of the distribution may demonstrate increased risk, this does not mean that PRS are useful for clinical purposes [16]. We therefore calculated the area under the curve (AUC) from receiver operating characteristic (ROC) curves as an additional check on any potential utility of PRS for alcohol use disorders (AUD) [17] using the *pROC* package in R[18]. ROC curves compare the sensitivity (true positive rate) and specificity (false positive rate) to identify an indicators ability to correctly classify an individual as having a disease/disorder or not.

Where the true positive rate (sensitivity) equals:

$$True\ Positive\ Rate\ = \frac{True\ Positives}{True\ Positives\ +\ False\ Negatives}$$

The false positive rate is therefore equal to:

$$False\ Positive\ Rate\ = \frac{False\ Positives}{False\ Positives\ +\ True\ Negatives}$$

We calculate these along the range of predicted values for models with only covariates and those with covariates + PRS, seen in Figure S3. As can be seen in the ROC curves, adding the PRS to the models with covariates is only slightly better. The AUC from the models with PRS are also provided. AUC provides an estimate of the probability of correctly classifying two randomly drawn individuals from the sample. An AUC of 0.5 indicates that a classifier does not provide any useful information in determining cases from controls. The AUC for the full models including PRS at each threshold suggest these models due a poor-to-moderate job at classifying individuals as having AUD or not.

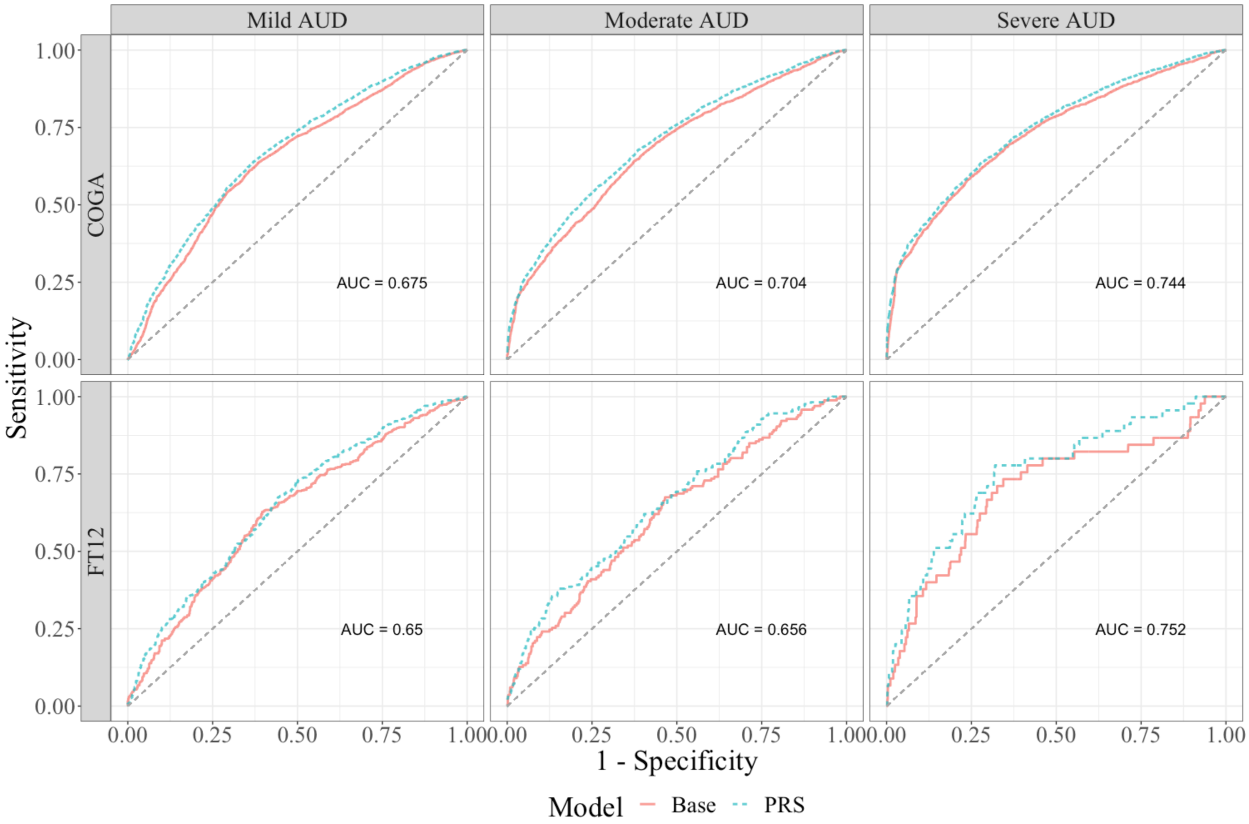


**Figure S4: ROC Curves for PRS Models**

#### 5. Restricting Age Range in COGA

Because the age ranges in COGA and FT12 varied so widely, we ran a series of sensitivity analyses to ensure that differences across samples were not the result of age differences. We reran all of the analyses in the manuscript restricting the COGA sample to those aged 18 to 30 years old.

While the results of predictive power, increase in prevalence of AUD across septiles, and joint influence of PRS are not substantively different from the results in the full sample, we see that the pattern at the extreme end of the PRS continuum did not follow a pattern of increasing risk. Instead, restricting to more extreme end of the PRS continuum resulted in a *smaller* OR relative to the less extreme thresholds.

**Table S3: Odds Ratios for Those at Extreme End of the PRS Continuum in COGA (ages 18-30)**

| Sample | Phenotype | Split | N Cases | OR | 95 % CI Low | 95 % CI High |
|---|---|---|---|---|---|---|
| | Mild AUD | 80% | 360 | 1.964 | 1.562 | 2.468 |
| COGA | Mild AUD | 90% | 182 | 1.680 | 1.247 | 2.262 |
| | Mild AUD | 95% | 90 | 1.488 | 0.992 | 2.232 |
| | Moderate AUD | 80% | 224 | 1.854 | 1.466 | 2.345 |
| COGA | Moderate AUD | 90% | 118 | 1.752 | 1.295 | 2.370 |
| | Moderate AUD | 95% | 60 | 1.584 | 1.054 | 2.379 |
| | Severe AUD | 80% | 131 | 1.621 | 1.236 | 2.125 |
| COGA | Severe AUD | 90% | 74 | 1.767 | 1.258 | 2.483 |
| | Severe AUD | 95% | 37 | 1.494 | 0.945 | 2.362 |

All models control for sex, age at last interview, and first 10 principal components. Models for COGA also included data collection site and genotyping array. N Cases = number of individuals who meet criteria for a given level of AUD and are in the top portion of the split.

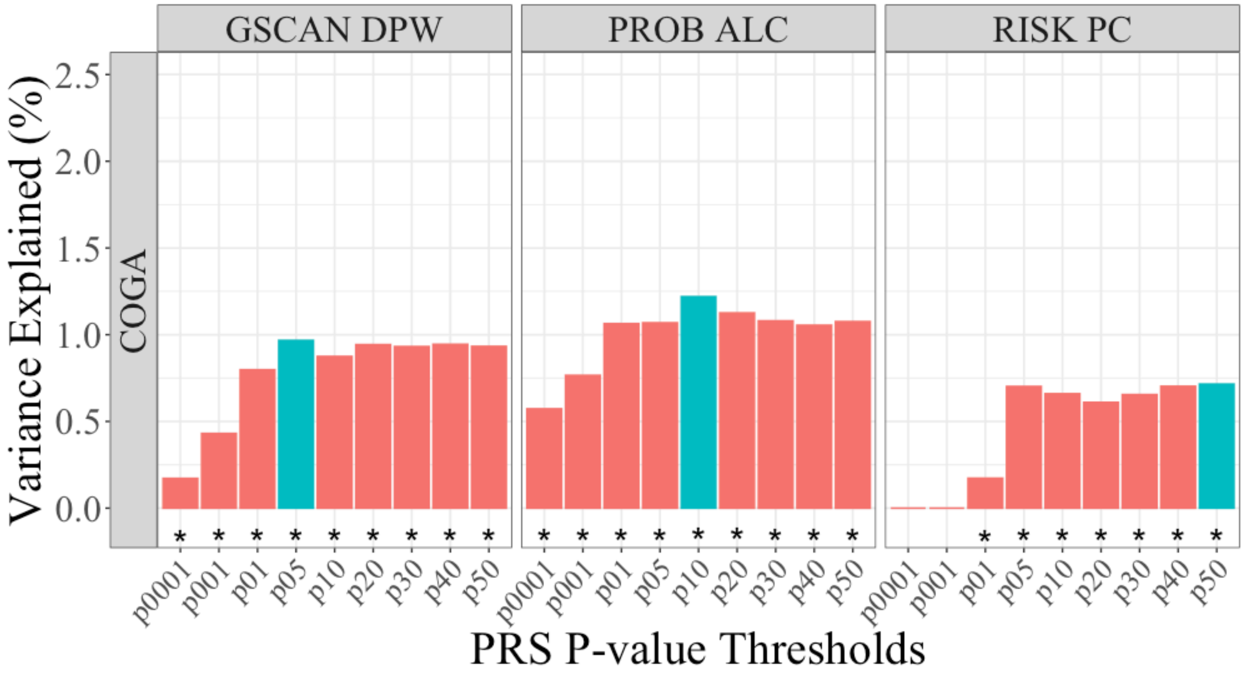


**Figure S5: Predictive Power of PRS**

Predictive power of PRS measured using pseudo-$R^2$ for linear mixed effects models [29]. Bars represent change in variance explained over models with age, sex, and first 10 ancestral principal components, genotyping array, and data collection site. Most predictive score outlined in blue. * p < .05, corrected for FDR of 5%

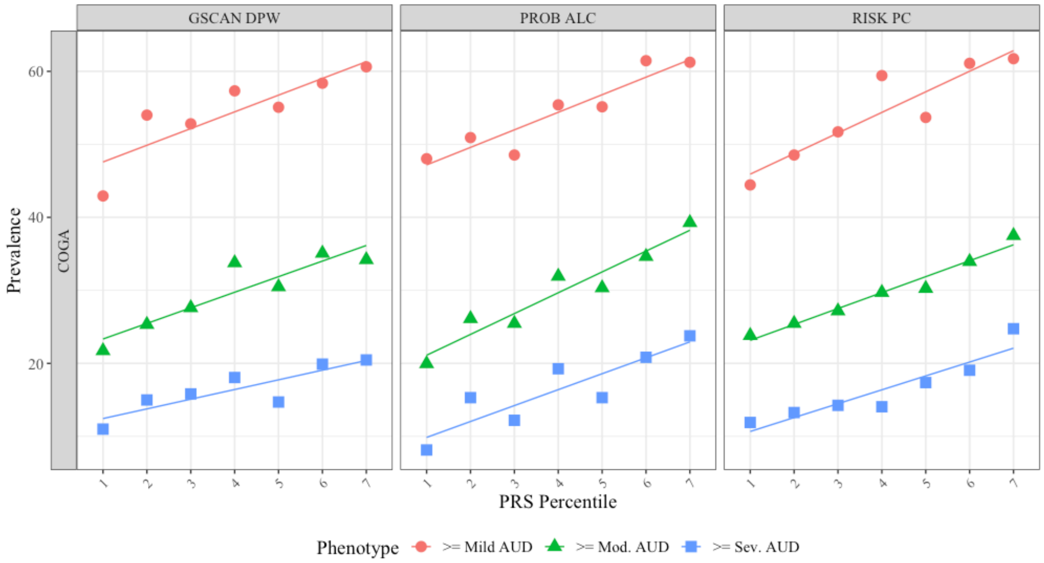


**Figure S6: Figure 7: Prevalence of AUD Across PRS Continuum**

Mean Prevalence of mild, moderate, and severe AUD in COGA (restricted to ages 18-30) across deciles for GSCAN DPW, PROB ALC, and RISK PC PRS. Adjusted for age, sex, first 10 ancestral principal components, genotyping array, and data collection site.

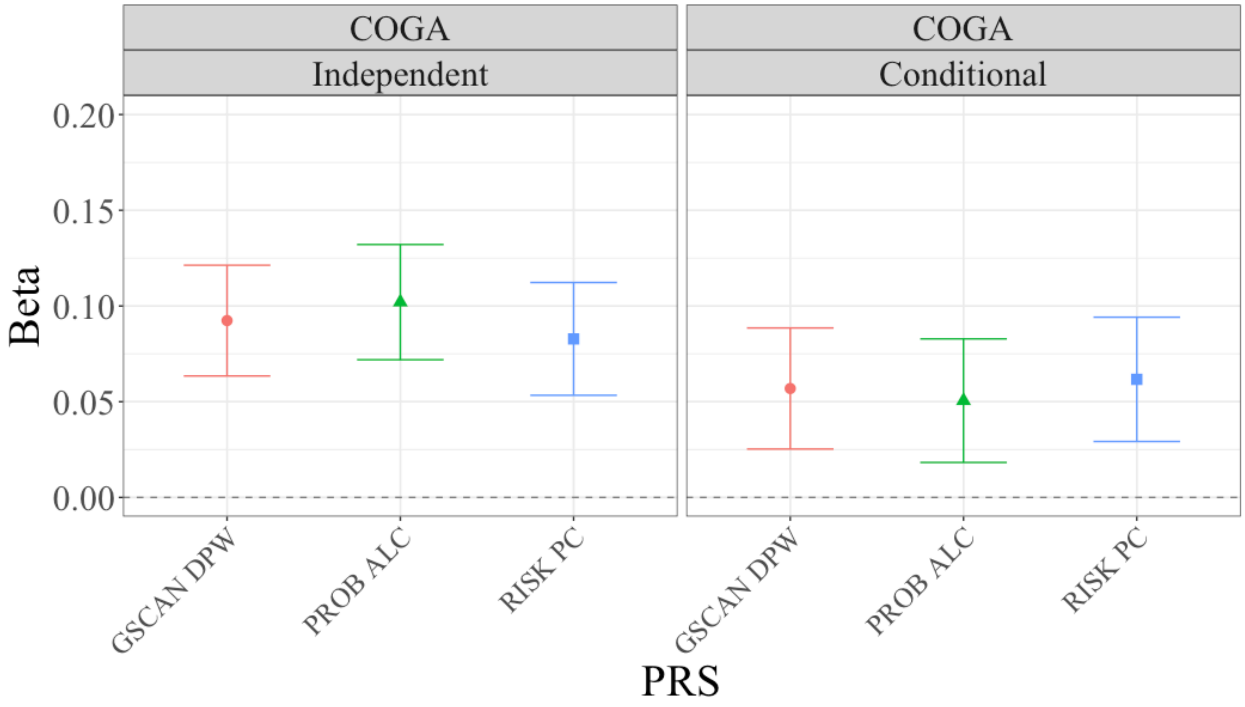


**Figure S8:** ***: Parameter Estimates for PRS in Independent and Joint Models***

Parameter estimates from linear mixed models for AUD symptoms regressed on GSCAN DPW, PROB ALC, and RISK PC PRS in COGA (age restricted). Independent = model with only corresponding PRS. Conditional = model with all PRS included. Adjusted for age, sex, first 10 ancestral principal components, genotyping array, and data collection site.

**References**

1. Edenberg HJ, Koller DL, Xuei X, Wetherill L, McClintick JN, Almasy L, . . . Foroud T. Genome-Wide Association Study of Alcohol Dependence Implicates a Region on Chromosome 11. Alcoholism: Clinical and Experimental Research. 2010;34(5):840-52. doi: 10.1111/j.1530-0277.2010.01156.x.
2. Wang JC, Foroud T, Hinrichs AL, Le NXH, Bertelsen S, Budde JP, . . . Goate AM. A genome-wide association study of alcohol-dependence symptom counts in extended pedigrees identifies C15orf53. Molecular psychiatry. 2012;18:1218. doi: 10.1038/mp.2012.143 https://www.nature.com/articles/mp2012143#supplementary-information.
3. Baurley JW, Edlund CK, Pardamean CI, Conti DV, Bergen AW. Smokescreen: a targeted genotyping array for addiction research. BMC Genomics. 2016;17(1):145. doi: 10.1186/s12864-016-2495-7.
4. O'Connell JR, Weeks DE. PedCheck: A Program for Identification of Genotype Incompatibilities in Linkage Analysis. The American Journal of Human Genetics. 1998;63(1):259-66. doi: https://doi.org/10.1086/301904.
5. Delaneau O, Howie B, Cox Anthony J, Zagury J-F, Marchini J. Haplotype Estimation Using Sequencing Reads. The American Journal of Human Genetics. 2013;93(4):687-96. doi: https://doi.org/10.1016/j.ajhg.2013.09.002.
6. Das S, Forer L, Schönherr S, Sidore C, Locke AE, Kwong A, . . . Fuchsberger C. Next-generation genotype imputation service and methods. Nature Genetics. 2016;48:1284. doi: 10.1038/ng.3656 https://www.nature.com/articles/ng.3656#supplementary-information.
7. Price AL, Patterson NJ, Plenge RM, Weinblatt ME, Shadick NA, Reich D. Principal components analysis corrects for stratification in genome-wide association studies. Nature Genetics. 2006;38:904. doi: 10.1038/ng1847 https://www.nature.com/articles/ng1847#supplementary-information.
8. Kaprio J. The Finnish Twin Cohort Study: an update. Twin Res Hum Genet. 2013;16(1):157-62. doi: 10.1017/thg.2012.142. PubMed PMID: 23298696; PMCID: PMC4493754.
9. The 1000 Genomes Project Consortium. A global reference for human genetic variation. Nature. 2015;526:68. doi: 10.1038/nature15393 https://www.nature.com/articles/nature15393#supplementary-information.
10. Delaneau O, Zagury J-F, Marchini J. Improved whole-chromosome phasing for disease and population genetic studies. Nature Methods. 2012;10:5. doi: 10.1038/nmeth.2307 https://www.nature.com/articles/nmeth.2307#supplementary-information.
11. Howie BN, Donnelly P, Marchini J. A Flexible and Accurate Genotype Imputation Method for the Next Generation of Genome-Wide Association Studies. PLOS Genetics. 2009;5(6):e1000529. doi: 10.1371/journal.pgen.1000529.
12. Bulik-Sullivan BK, Loh PR, Finucane HK, Ripke S, Yang J, Schizophrenia Working Group of the Psychiatric Genomics C, . . . Neale BM. LD Score regression distinguishes confounding from polygenicity in genome-wide association studies. Nat Genet. 2015;47(3):291-5. doi: 10.1038/ng.3211. PubMed PMID: 25642630; PMCID: PMC4495769.
13. Willer CJ, Li Y, Abecasis GR. METAL: fast and efficient meta-analysis of genomewide association scans. Bioinformatics. 2010;26(17):2190-1. doi: 10.1093/bioinformatics/btq340. PubMed PMID: 20616382; PMCID: PMC2922887.
14. Khera AV, Chaffin M, Aragam KG, Haas ME, Roselli C, Choi SH, . . . Kathiresan S. Genome-wide polygenic scores for common diseases identify individuals with risk equivalent to monogenic mutations. Nature Genetics. 2018;50(9):1219-24. doi: 10.1038/s41588-018-0183-z.
15. Khera AV, Chaffin M, Wade KH, Zahid S, Brancale J, Xia R, . . . Kathiresan S. Polygenic Prediction of Weight and Obesity Trajectories from Birth to Adulthood. Cell. 2019;177(3):587-96 e9. doi: 10.1016/j.cell.2019.03.028. PubMed PMID: 31002795.
16. Wald NJ, Old R. The illusion of polygenic disease risk prediction. Genetics in Medicine. 2019. doi: 10.1038/s41436-018-0418-5.
17. Steyerberg EW, Vickers AJ, Cook NR, Gerds T, Gonen M, Obuchowski N, . . . Kattan MW. Assessing the performance of prediction models: a framework for traditional and novel measures. Epidemiology. 2010;21(1):128-38. doi: 10.1097/EDE.0b013e3181c30fb2. PubMed PMID: 20010215; PMCID: PMC3575184.

18. Robin X, Turck N, Hainard A, Tiberti N, Lisacek F, Sanchez J-C, Müller M. pROC: an open-source package for R and S+ to analyze and compare ROC curves. BMC Bioinformatics. 2011;12(1):77. doi: 10.1186/1471-2105-12-77.
